# TAX1BP1 recruits ATG9 vesicles through SCAMP3 binding

**DOI:** 10.1101/2023.08.18.553817

**Authors:** Yutaro Hama, Yoshitaka Kurikawa, Takahide Matsui, Noboru Mizushima, Hayashi Yamamoto

**Author notes:** Correspondence: Hayashi Yamamoto, Noboru Mizushima.

## Abstract

Macroautophagy is a cellular process that delivers cytoplasmic material to lysosomes for degradation via autophagosomes. It often involves the selective degradation of ubiquitinated proteins. During selective macroautophagy, five ubiquitin-binding adaptors, p62, NBR1, OPTN, NDP52, and TAX1BP1, form biomolecular condensates with ubiquitinated proteins and recruit ATG9 vesicles, which serve as the initial membrane source required for autophagosome formation. However, the molecular details underlying the cargo/adaptor-dependent recruitment of ATG9 vesicles remain unclear. Here, we show that ATG9 vesicles are recruited by three cargo adaptors: TAX1BP1, NBR1, and OPTN. We also find that ATG9A itself is not the determinant for recruitment by these cargo adaptors, and that TAX1BP1-dependent ATG9 vesicle recruitment is mediated by SCAMP3, a transmembrane protein on the ATG9 vesicles, through binding to the coiled-coil 1 domain of TAX1BP1. These findings provide mechanistic insights into the cargo/adaptor-dependent assembly of ATG9 vesicles in mammals.

## Introduction

Macroautophagy (hereafter referred to as autophagy) is an intracellular degradation system that sequesters cytoplasmic materials in autophagosomes nonselectively or selectively and delivers them to lysosomes or vacuoles for degradation (Mizushima and Komatsu 2011; Yamamoto et al. 2023; Yamamoto and Matsui 2023). Recently, selective autophagy has attracted much attention because of its involvement in human diseases, including neurodegenerative diseases (Xu et al. 2021; Yamamoto et al. 2023). One of the major targets of selective autophagy in mammals is ubiquitinated proteins, which are recognized by the selective cargo adaptors (also called receptors) p62 (also known as SQSTM1), NBR1, OPTN, NDP52 (also known as CALCOCO2), and TAX1BP1, all of which possess ubiquitin-binding domains. These five adaptors also possess the LC3-interacting region and thus bind to the LC3 family proteins localized on the autophagosomal membranes, thereby mediating selective incorporation of ubiquitinated proteins and organelles into autophagosomes (Johansen and Lamark 2020).

Autophagosome formation requires hierarchical recruitment and cooperative functions of core autophagy-related (ATG) proteins (Nakatogawa 2020; Yamamoto et al. 2023), the first step of which involves assembly of the ULK complex consisting of ULK1 (or ULK2), ATG13, ATG101, and FIP200 (also known as RB1CC1), and recruitment of ATG9 vesicles, which serve as the initial membrane source for autophagosome formation (Yamamoto et al. 2012; Sawa-Makarska et al. 2020). The ULK complex and ATG9 vesicles are recruited to autophagosome formation sites not only in a starvation-induced manner, but also in a cargo-driven manner. In the cargo-driven pathway (Fig. 1A), FIP200, a central subunit of the ULK complex, directly interacts with the cargo adaptors p62, TAX1BP1, and NDP52 (Ravenhill et al. 2019; Turco et al. 2019; Vargas et al. 2019; Ohnstad et al. 2020; Turco et al. 2021). FIP200 binds to p62 via its C-terminal Claw domain (Turco et al. 2019), resulting in the recruitment of the ULK complex to ubiquitin- and p62-double-positive structures, namely the p62 condensates (Kishi-Itakura et al. 2014). p62 condensates are characterized as biomolecular condensates containing ubiquitinated proteins and five cargo adaptors: p62, NBR1, OPTN, NDP52, and TAX1BP1 (Kirkin et al. 2009; Cemma et al. 2011; Wong and Holzbaur 2014; Tumbarello et al. 2015; Wurzer et al. 2015; Sun et al. 2018). p62 is the main driver to produce the p62 condensates, and the recruitment of the ULK complex to these condensates leads to the efficient degradation of ubiquitinated cargos. TAX1BP1, another component of the p62 condensates, facilitates the recruitment of the ULK complex by binding to FIP200 (Turco et al. 2021). FIP200 also directly interacts with the skeletal muscle and kidney-enriched inositol phosphatase carboxyl homology (SKICH) domain of NDP52, resulting in the recruitment of the ULK complex to ubiquitinated mitochondria during mitophagy and to invading bacteria during xenophagy (Ravenhill et al. 2019; Vargas et al. 2019; Shi et al. 2020) (Fig. 1A).

**Figure 1.**
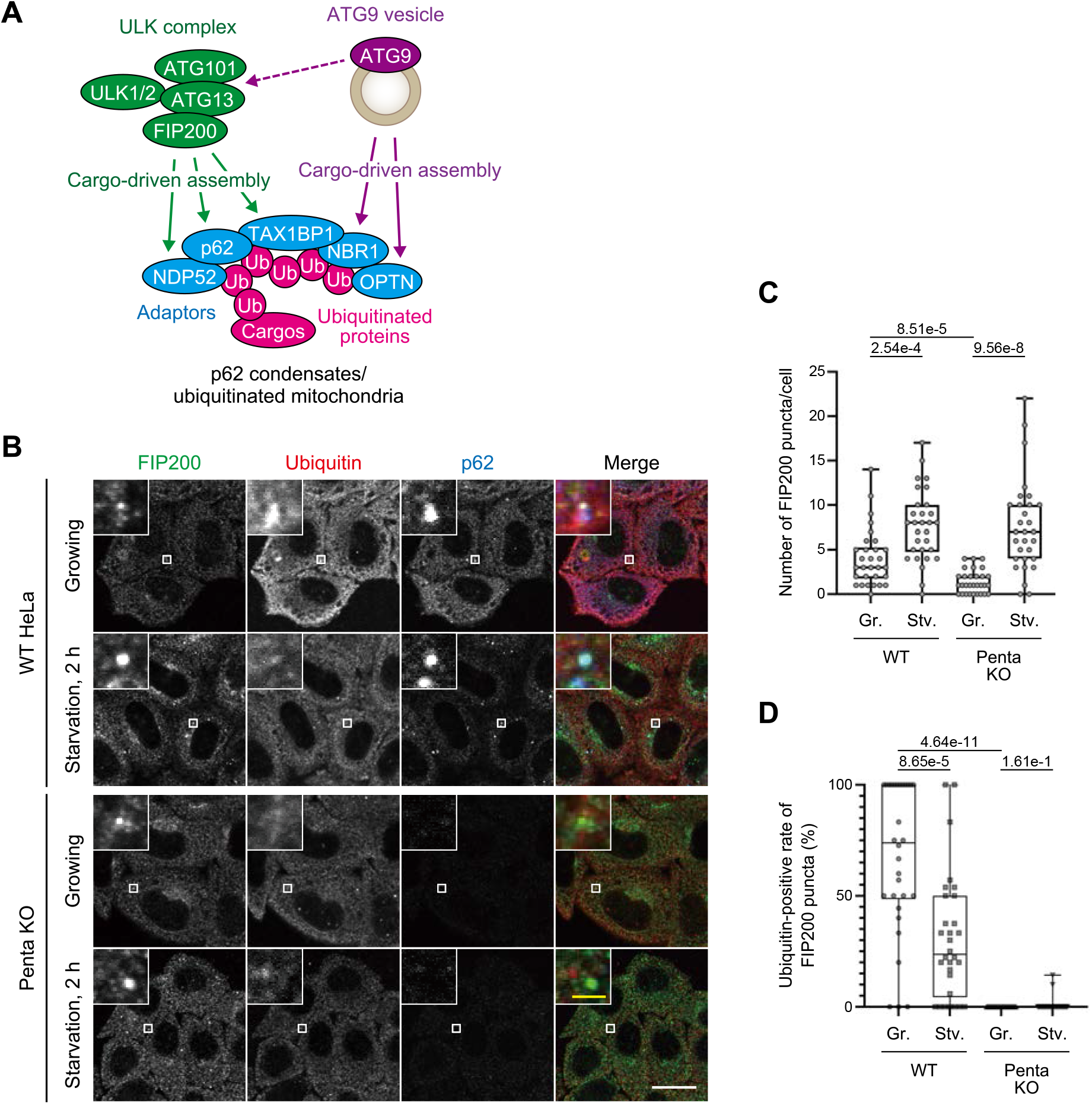
Autophagy adaptors are required for cargo-dependent but not starvation-induced assembly of the ULK complex. **(A)** Schematic representation of cargo-driven assembly of early ATG proteins such as the ULK complex and ATG9 vesicles during selective autophagy against p62 condensates and ubiquitinated mitochondria. The cargo-driven assembly is indicated by solid arrows and the ULK complex-dependent ATG9 vesicle recruitment is indicated by a dashed arrow. **(B)** Starvation-induced assembly of the ULK complex is distinguished by deletion of five cargo adaptors. WT and Penta KO (deletion of p62, NBR1, NDP52, OPTN, and TAX1BP1) HeLa cells were starved for 2 h and immunostained with anti-ATG9A, anti-ubiquitin, and anti-p62 antibodies. **(C)** The number of FIP200 puncta per cell in (B) was counted (30 cells each). Solid bars indicate the means, boxes the interquartile ranges, and whiskers the 25th to 75th percentiles. Differences were statistically analyzed by the Mann-Whitney test. **(D)** The ratios of ubiquitin-positive FIP200 puncta to total FIP200 puncta per cell in (B) were calculated (30 cells each). Solid bars indicate the means, boxes the interquartile ranges, and whiskers the 25th to 75th percentiles. Differences were statistically analyzed by the Mann-Whitney test.

In contrast, how ATG9 vesicles are recruited in a cargo-dependent manner remains poorly understood. A recent study showed that during Parkin-mediated mitophagy, the ULK complex recruits ATG9 vesicles through direct interaction between ATG9A and the ATG13–ATG101 subcomplex in the ULK complex (Kannangara et al. 2021; Ren et al. 2023). In addition, ATG9 vesicles are also recruited to the p62 condensates in the absence of the ULK complex (Kishi-Itakura et al. 2014), and during mitophagy, ATG9 vesicles localize to depolarized mitochondria independently of the ULK complex (Itakura et al. 2012). Several studies have reported that ATG9 vesicles are recruited by two of the five cargo adaptors, OPTN, which localizes to ubiquitinated mitochondria (Yamano et al. 2020), and NBR1, which is included in the p62 condensates (Sánchez-Martín et al. 2020). However, it remains unclear whether other cargo adaptors are involved in the cargo-dependent recruitment of ATG9 vesicles.

In this study, we investigated cargo/adaptor-dependent and starvation-induced recruitment of the ULK complex and ATG9 vesicles using Penta knockout (KO) cells, which lack all five cargo adaptors. Fluorescence microscopy revealed that ATG9 vesicles are recruited by TAX1BP1, as well as OPTN and NBR1, independently of the ULK complex. We identified SCAMP3 as a protein on the ATG9 vesicles that binds to TAX1BP1. SCAMP3 binding is responsible for the TAX1BP1-dependent ATG9 vesicle recruitment. These findings facilitate a better understanding of the cargo/adapter-dependent assembly of ATG9 vesicles in mammals.

## Results

### Autophagy adaptors are required for cargo-dependent but not starvation-induced assembly of the ULK complex

We first determined how cargo-dependent and starvation-induced assembly of the ULK complex occurs. In wild-type (WT) HeLa cells, FIP200, a central subunit of the ULK complex, formed punctate structures adjacent to ubiquitin-and p62-double-positive structures under growing conditions (Fig. 1B, 1C), as previously reported (Kishi-Itakura et al. 2014), which represent cargo (p62 condensate)-driven recruitment of the ULK complex. In contrast, in Penta KO cells, in which all five cargo adaptors (p62, NBR1, OPTN, NDP52, and TAX1BP1) were deleted (Lazarou et al. 2015), FIP200 puncta were significantly reduced in number (Fig. 1B, 1C) and exhibited less colocalization with ubiquitin-positive condensates under growing conditions (Fig. 1B, 1D). Expression of NDP52, NBR1, or TAX1BP1, but not p62 or OPTN, restored the ULK complex assembly to ubiquitin-positive condensates (Fig. S1A, S1B), which suggests that cargo-dependent recruitment of the ULK complex requires the presence of NDP52, NBR1, or TAX1BP1. In response to starvation, the number of FIP200 puncta was significantly increased even in Penta KO cells (Fig. 1B, 1C), and the ubiquitin– FIP200 colocalization rate was decreased as compared with that observed in WT cells (Fig. 1B, 1D), representing cargo- and adaptor-independent assembly of the ULK complex. These results suggest the necessity of the autophagy adaptors in cargo-dependent but not starvation-induced assembly of the ULK complex.

### ATG9 vesicles colocalize with the ULK complex dependent on the HORMA domain of ATG13 in the absence of the adaptors

We next examined the recruitment of ATG9 vesicles. ATG9 vesicles are known to colocalize with the ULK complex during starvation (Koyama-Honda et al. 2013). This co-localization occurred even in Penta KO cells (Fig. 2A, 2B), which indicates that starvation-induced recruitment of ATG9 vesicles does not require adaptors. This recruitment depended on the HORMA domain of ATG13, indicating that the interaction between ATG9A and the ATG13–ATG101 subcomplex is required for not only selective autophagy (Ren et al. 2023) but also starvation-induced non-selective autophagy (Fig. 2A, 2B). These results suggest that the starvation-induced recruitment of ATG9 vesicles depends on the binding to ATG13–ATG101 in the absence of the adaptors.

**Figure 2.**
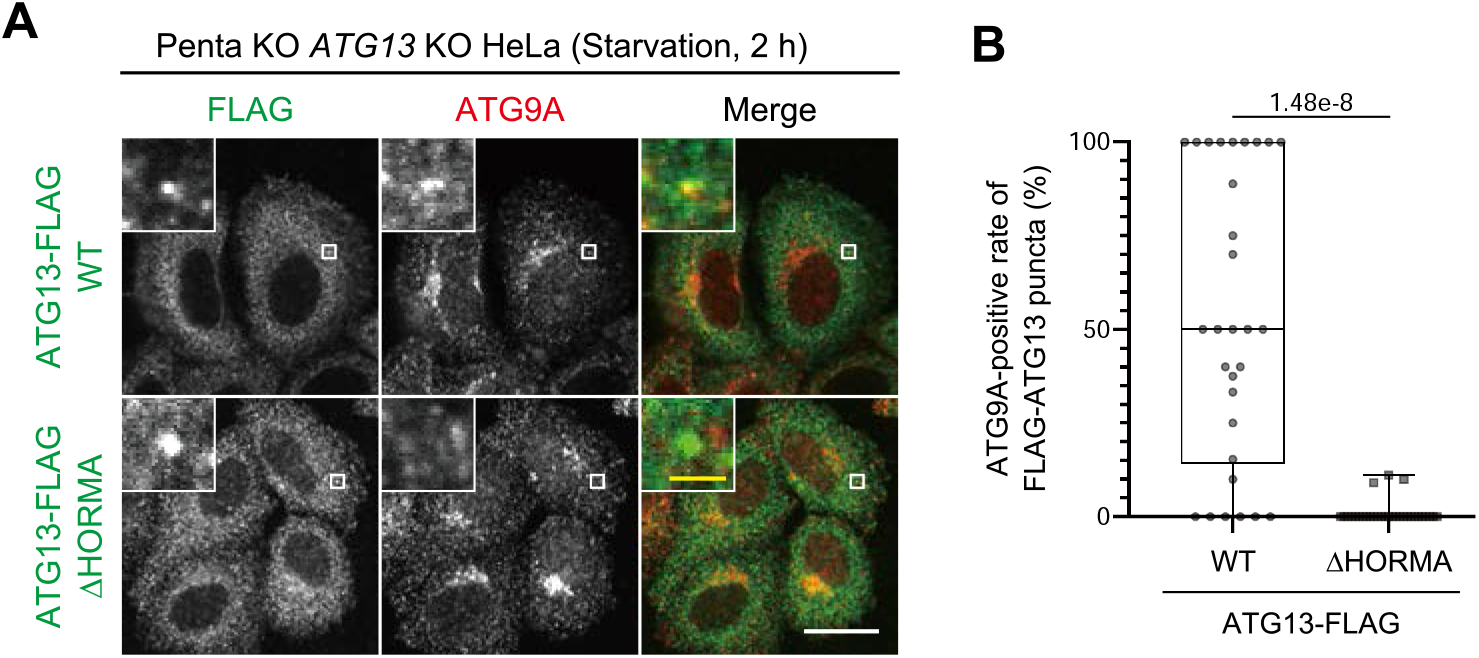
ATG9 vesicles colocalize with the ULK complex dependent on the HORMA domain of ATG13 in the absence of adaptors. **(A)** ATG9 vesicles localize to the starvation-induced assembly of the ULK complex in an ATG13^HORMA^-dependent manner. Penta KO *ATG13* KO HeLa cells stably expressing WT or ΔHORMA mutant of ATG13-FLAG were starved for 2 h and immunostained with anti-FLAG and anti-ATG9A antibodies. **(B)** The ratios of ATG9A-positive ATG13-FLAG puncta to total ATG13-FLAG puncta per cell were calculated (30 cells each) as in Figure 1D. White scale bars represent 20 μm and yellow scale bars represent 2 μm.

### TAX1BP1, NBR1, and OPTN recruit ATG9 vesicles during cargo-dependent autophagy

We next examined cargo-dependent recruitment of ATG9 vesicles. In WT cells, a portion of ATG9A colocalized with the ubiquitin–p62 condensates under growing conditions (Fig. 3A, 3B). The colocalization rate was significantly increased in the absence of FIP200 (Fig. 3A, 3B), which is in agreement with previous studies showing that ATG9 vesicles are recruited to the p62 condensates in a ULK complex-independent manner (Kishi-Itakura et al. 2014). In Penta KO cells, ubiquitin-positive condensates were still observed, but ATG9A did not localize to these condensates, even when FIP200 was simultaneously deleted (Penta KO *FIP200* KO cells) (Fig. 3A, 3B). These results indicate that ATG9 vesicles are recruited to the ubiquitin–p62 condensates in a manner dependent on the cargo adaptors, but not on the ULK complex.

**Figure 3.**
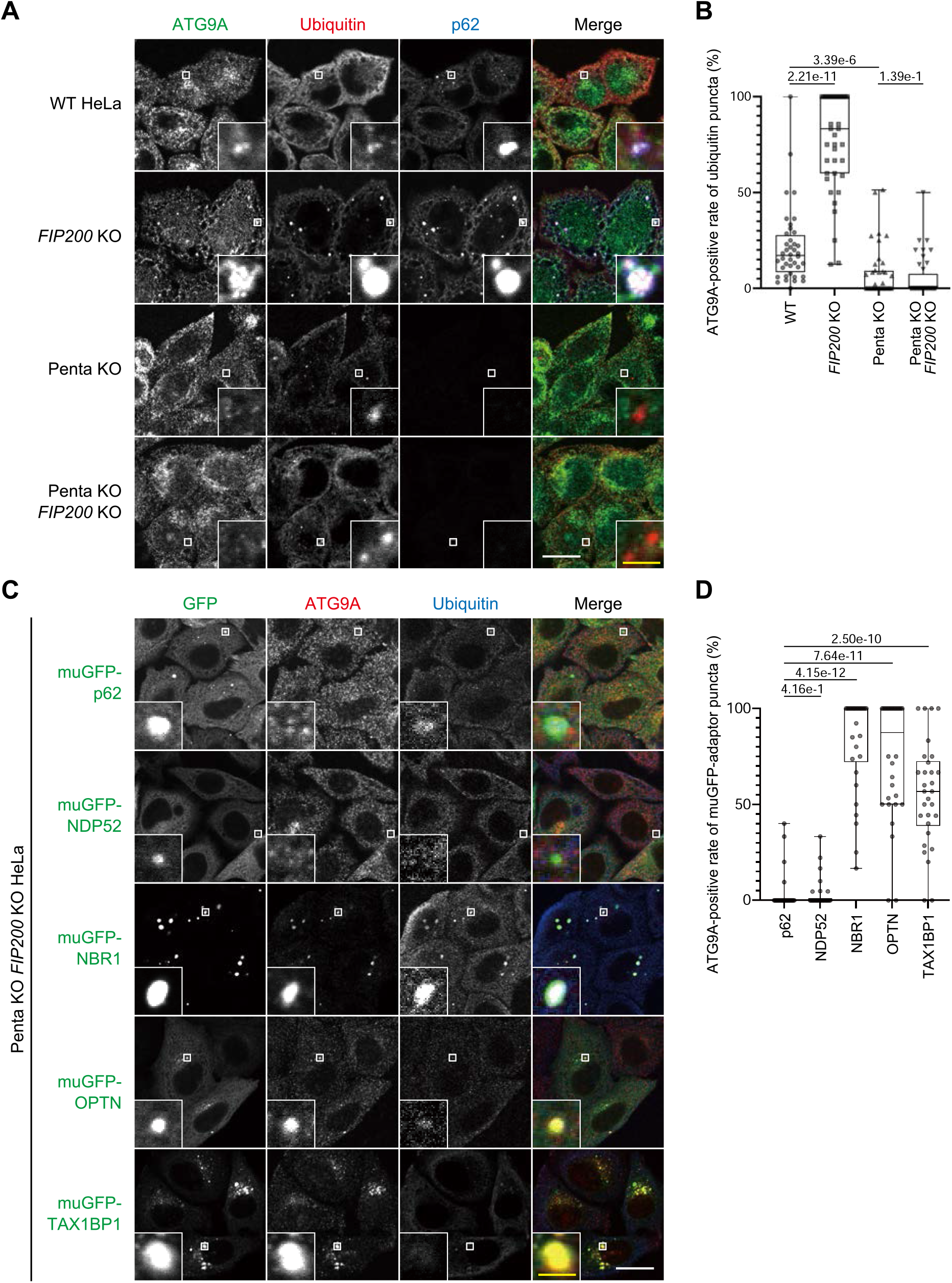
TAX1BP1, NBR1, and OPTN recruit ATG9 vesicles independently of each other and of the ULK complex. **(A)** ATG9 vesicles do not localize to ubiquitin-positive puncta in the absence of five cargo adaptors p62, NBR1, OPTN, NDP52, and TAX1BP1. WT, *FIP200* KO, Penta KO, and Penta KO *FIP200* KO HeLa cells were immunostained with anti-ATG9A, anti-ubiquitin, and anti-p62 antibodies. **(B)** The ratios of ATG9A-positive ubiquitin puncta to total ubiquitin puncta per cell in (A) were calculated (40 cells each) as in Figure 1D. **(C)** ATG9 vesicles localize to NBR1, OPTN, and TAX1BP1 puncta, but not to p62 and NDP52 puncta. Penta KO *FIP200* KO HeLa cells stably expressing muGFP-tagged p62, NBR1, OPTN, NDP52, or TAX1BP1 were immunostained with anti-GFP, anti-ATG9A, and anti-ubiquitin antibodies. **(D)** The ratios of ATG9A-positive GFP puncta to total GFP puncta per cell in (C) were calculated (40 cells each) as in Figure 1D. White scale bars represent 20 μm and yellow scale bars represent 2 μm.

We next investigated which adaptors are required to recruit ATG9 vesicles. The expression of any of the monomeric ultra-stable green fluorescent protein (muGFP)-tagged cargo adaptors produced punctate structures in Penta KO *FIP200* KO cells (Fig. 3C). Of these, NBR1, OPTN, and TAX1BP1 puncta were positive for ATG9A, whereas p62 and NDP52 puncta were negative for ATG9A (Fig. 3C, 3D), suggesting that NBR1, OPTN, and TAX1BP1 recruit ATG9 vesicles independently of other adaptors.

### The coiled-coil 1 domain of TAX1BP1 is required for ATG9 vesicle recruitment

Because NBR1 and OPTN have been reported to be involved in ATG9 vesicle recruitment (Sánchez-Martín et al. 2020; Yamano et al. 2020), we focused on TAX1BP1. TAX1BP1 consists of six domains: the SKICH, noncanonical LC3C-interacting region (CLIR), coiled-coil 1 (CC1), CC2, CC3, and ubiquitin-binding zinc finger (UBZ) domains (Fig. 4A). To determine which domain is required for ATG9 vesicle recruitment, we constructed domain deletion mutants of muGFP-TAX1BP1 (ΔSKICH, ΔCLIR, ΔCC1, ΔCC2, ΔCC3, and ΔUBZ) (Fig. 4A).

**Figure 4.**
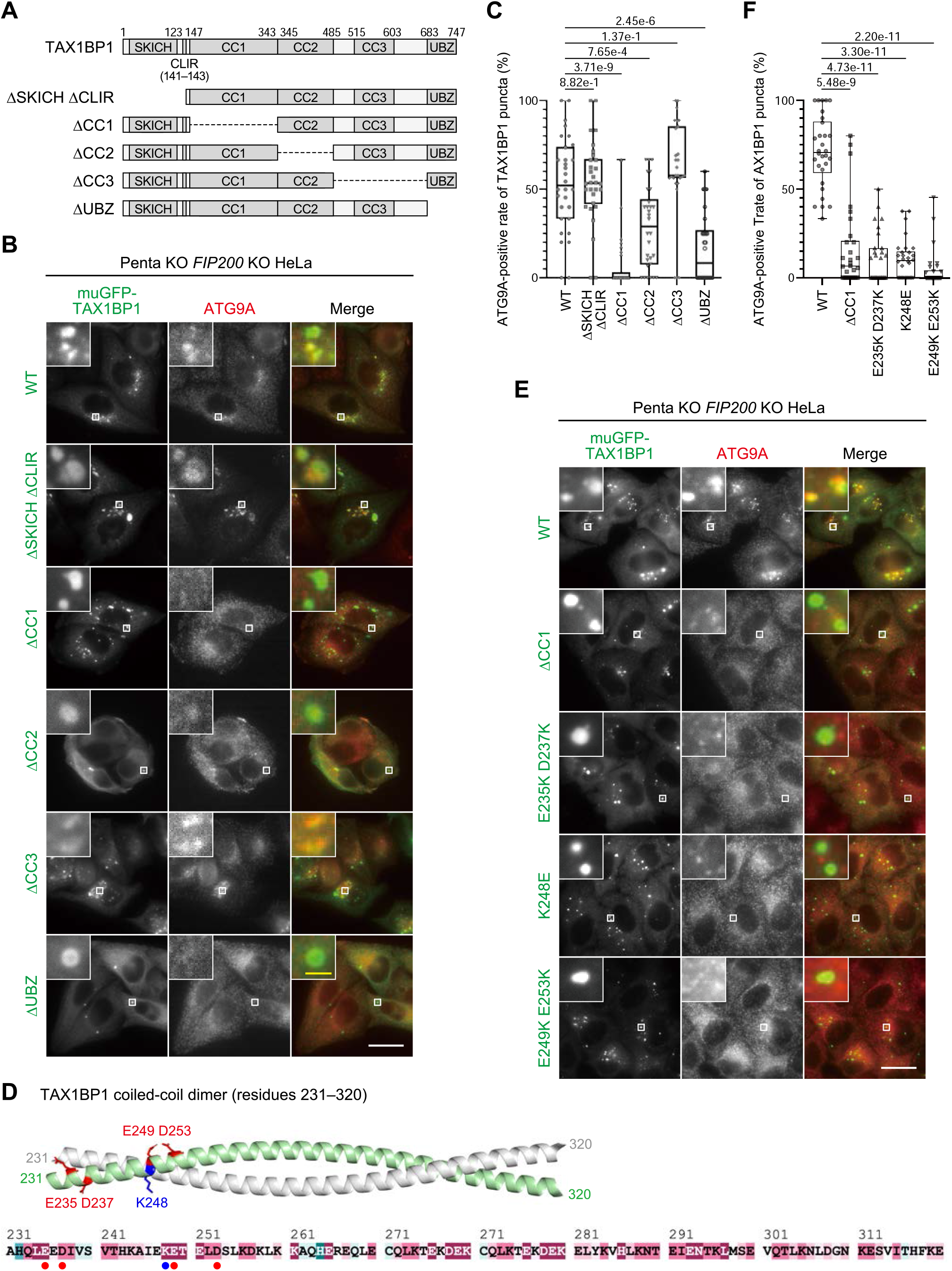
TAX1BP1 recruits ATG9 vesicles via the CC1 domain. **(A)** Truncated mutants of TAX1BP1. TAX1BP1 consists of SKICH, CLIR, CC1, CC2, CC3, and UBZ domains. **(B)** The CC1 domain of TAX1BP1 (residues 147–343) is required for ATG9 vesicle recruitment. Penta KO *FIP200* KO HeLa cells stably expressing WT or mutants (ΔSKICH, ΔCLIR, ΔCC1, ΔCC2, ΔCC3, or ΔUBZ) of muGFP-TAX1BP1 were immunostained with anti-GFP, anti-ATG9A, and anti-ubiquitin antibodies. **(C)** The ratios of ATG9A-positive GFP puncta to total GFP puncta per cell in (B) were calculated (30 cells each) as in Figure 1D. **(D)** The structure of TAX1BP1^CC1^ (residues 147–343) as predicted by AlphaFold-Multimer and the residues 231–320 are shown (top). The amino-acid conservation of TAX1BP1^CC1^ analyzed by the ConSurf server is shown with a heat map (bottom). Mutation points are indicated in red and blue. **(E)** Charged residues of the CC1 domain in TAX1BP1 are involved in ATG9 vesicle recruitment. Penta KO *FIP200* KO HeLa cells stably expressing WT or mutants (ΔCC1, E235K D237K, K248E, or E249K E253K) of muGFP-TAX1BP1 were immunostained with anti-GFP and anti-ATG9A antibodies. **(F)** The ratios of ATG9A-positive GFP puncta to total GFP puncta per cell in (E) were calculated (30 cells each) as in Figure 1D. White scale bars represent 20 μm and yellow scale bars represent 2 μm.

Fluorescence microscopy showed that all muGFP-TAX1BP1 mutants were observed as punctate structures in Penta KO *FIP200* KO cells (Fig. 4B). However, when the CC2 or UBZ domain was deleted, ATG9A only partially localized to the muGFP-TAX1BP1 puncta, and when the CC1 domain was deleted, ATG9A did not localize to the muGFP-TAX1BP1 puncta (Fig. 4B, 4C). Therefore, we introduced several mutations into the CC1 domain of TAX1BP1 (TAX1BP1^CC1^; residues 147– 343). Structural prediction by AlphaFold-Multimer (Evans et al. 2022; Mirdita et al. 2022) showed that TAX1BP1^CC1^ forms a coiled-coil homodimer, and amino-acid conservation analysis by ConSurf (Yariv et al. 2023) showed that residues 231– 320 are highly conserved. Therefore, we introduced mutations into the conserved charged residues facing outwards (E235K D237K, K248E, or E249K D253K) (Fig. 4D). All of the muGFP-TAX1BP1 mutants formed punctate structures in Penta KO *FIP200* KO cells, but ATG9A recruitment to these structures was significantly impaired in all mutants (Fig. 4E), suggesting that ATG9 vesicle recruitment depends on the surface charge of TAX1BP1^CC1^.

### TAX1BP1 interacts with SCAMP3, a transmembrane protein on the ATG9 vesicles

To elucidate the molecular mechanism by which TAX1BP1 recruits ATG9 vesicles, we first tried to detect the interaction between ATG9A and either WT or the K248E mutant of TAX1BP1^CC1^, but neither interaction was detected (Fig. 5A). Therefore, to identify the ATG9 vesicle component(s) that interacts with TAX1BP1^CC1^, we performed liquid chromatography–tandem mass spectrometry (LC–MS/MS) analysis of the FLAG-TAX1BP1^CC1^ immunoprecipitates. SCAMP3 was identified as an approximately three-fold enriched protein copurified with WT FLAG-TAX1BP1^CC1^ as compared with the K248E mutant, whereas ATG9A and ATG9B were not detected (Fig. 5B and Table S1). Co-immunoprecipitation also showed the interaction between TAX1BP1^CC1^ and SCAMP3; FLAG-TAX1BP1^CC1^ precipitated SCAMP3 under detergent-solubilized conditions (Fig. 5A, 5C). SCAMP3 is a tetra-spanning membrane protein localized to extracellular vesicles, the Golgi apparatus, and endosomes (Brand et al. 1991; Aoh et al. 2009), and has been identified in the proteome of ATG9 vesicles (Kakuta et al. 2017). We also confirmed that ATG9 vesicles isolated from HeLa cells contained SCAMP3 as well as RAB1A, a known component of the ATG9 vesicles (Fig. 5D). In addition, SCAMP3-mRFP localized to p62 condensates in WT cells, presumably as a component of ATG9 vesicles, and, like ATG9A, massively accumulated in p62 condensates in *FIP200* KO cells (Fig. 5E, 5F). Taken together, we conclude that SCAMP3 is a component of the ATG9 vesicles that interacts with TAX1BP1^CC1^.

**Figure 5.**
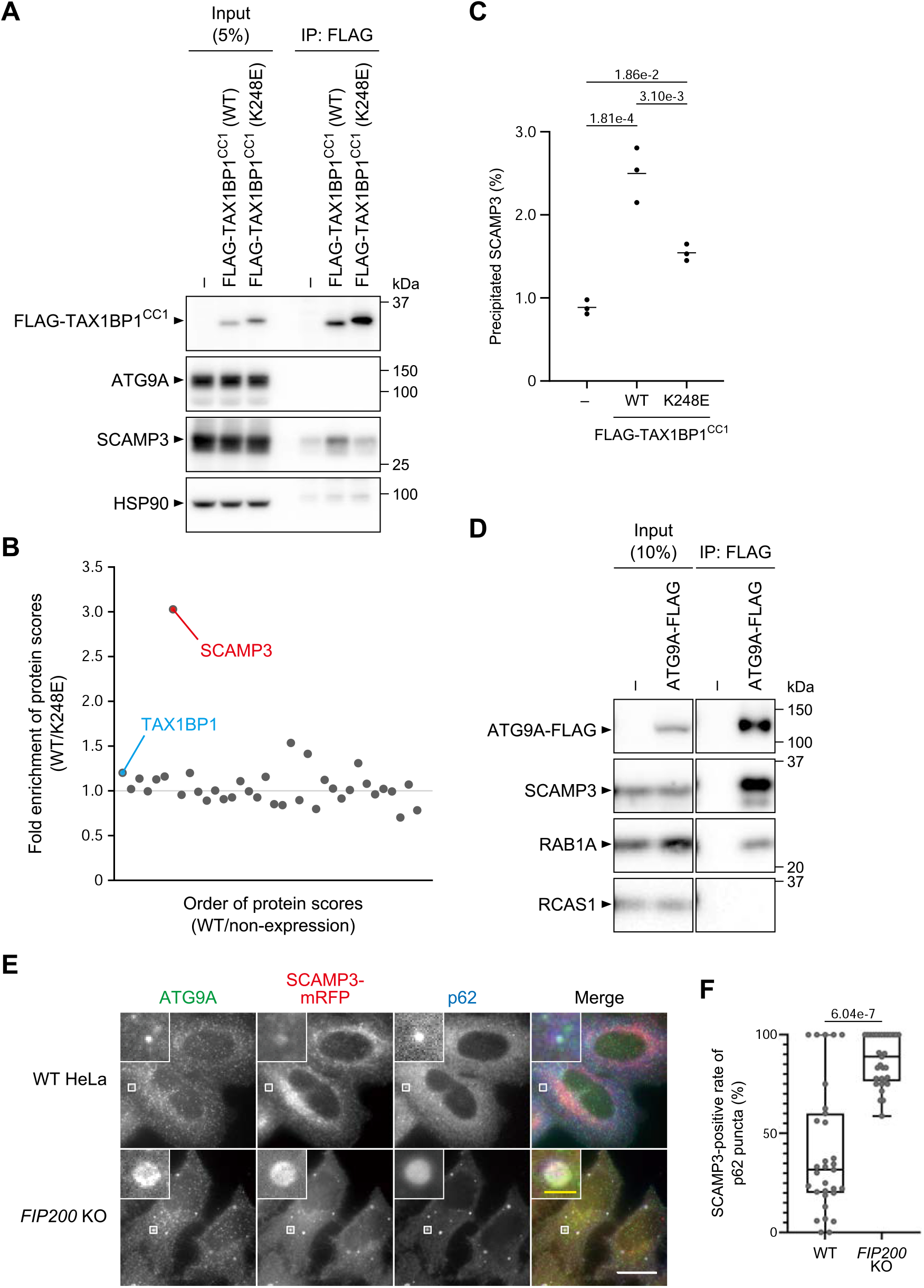
SCAMP3, an ATG9 vesicle component, interacts with TAX1BP1^CC1^. **(A)** TAX1BP1^CC1^ precipitates SCAMP3, but not ATG9A. Penta KO *FIP200* KO HeLa cells with or without stable expression of FLAG-TAX1BP1^CC1^ (WT or K248E) were solubilized with 0.2% n-octyl-β-D-dodecyl maltoside (DDM) and subjected to immunoprecipitation with anti-FLAG beads. The immunoprecipitates were detected with anti-FLAG, anti-ATG9A, anti-SCAMP3, and anti-HSP90 antibodies. **(B)** LC–MS/MS results of the TAX1BP1^CC1^ interactome. Immunoprecipitation with anti-FLAG beads was performed as in (A). The immunoprecipitates were labeled by dimethyl labeling to FLAG-TAX1BP1^CC1^ (WT) as heavy (H), FLAG-TAX1BP1 (K248E) as medium (M), and non-expression of TAX1BP1 as light (L) and analyzed by LC–MS/MS. The ratios of FLAG-TAX1BP1^CC1^ WT to FLAG-TAX1BP1^CC1^ K248E are plotted ([H/L]/[M/L]) for the proteins detected more than 1.5-fold by FLAG-TAX1BP1^CC1^ WT compared with non-expression (H/L>1.5). The x-axis shows the order of H/L, and the y-axis shows [H/L]/[M/L]. **(C)** Percentages of precipitated SCAMP3 in (A) were calculated. Solid bars indicate the means and dots indicate the data from three independent experiments. Differences were statistically analyzed by the Tukey-Kramer test. **(D)** SCAMP3 is copurified with ATG9 vesicles. WT HeLa cells stably expressing ATG9A-FLAG were subjected to immunopurification under non-detergent conditions, and the proteins copurified with ATG9 vesicles were detected with anti-FLAG, anti-SCAMP3, anti-RAB1A, and anti-RCAS1 antibodies. **(E)** SCAMP3 localizes to the p62 condensates. WT and *FIP200* KO HeLa cells stably expressing SCAMP3-mRFP were immunostained with anti-ATG9A and anti-p62 antibodies. **(F)** The ratios of SCAMP3-mRFP-positive p62 puncta to total p62 puncta per cell in (E) were calculated (30 cells each) in Figure 1D. White scale bars represent 20 μm and yellow scale bars represent 2 μm.

### SCAMP3 is responsible for ATG9 vesicle recruitment by TAX1BP1

To investigate the role of SCAMP3 in ATG9 vesicle recruitment to p62 condensates, we generated *SCAMP3 FIP200* double KO HeLa cells (Fig. S2). ATG9A localization to p62 condensates was partially but significantly reduced by deletion of the *SCAMP3* gene (Fig. 6A, 6B). Note that the *FIP200* KO background was used in the experiments in Figure 6 to investigate the ULK complex-independent mechanisms. Because ATG9A localization to p62 condensates was still partially observed even without SCAMP3, we hypothesized that TAX1BP1, NBR1, and OPTN did not require SCAMP3 to recruit ATG9 vesicles. Therefore, we conducted an analysis in which the *SCAMP3* gene was deleted in the Penta KO background, and muGFP-tagged TAX1BP1, NBR1, or OPTN was expressed individually. Fluorescence microscopy showed that *SCAMP3* KO significantly reduced the localization of ATG9A to muGFP-TAX1BP1 condensates, but not to muGFP-NBR1 or muGFP-OPTN condensates (Fig. 6C, 6D). These results indicate that SCAMP3 is responsible for the recruitment of ATG9 vesicles by TAX1BP1, but not by NBR1 or OPTN.

**Figure 6.**
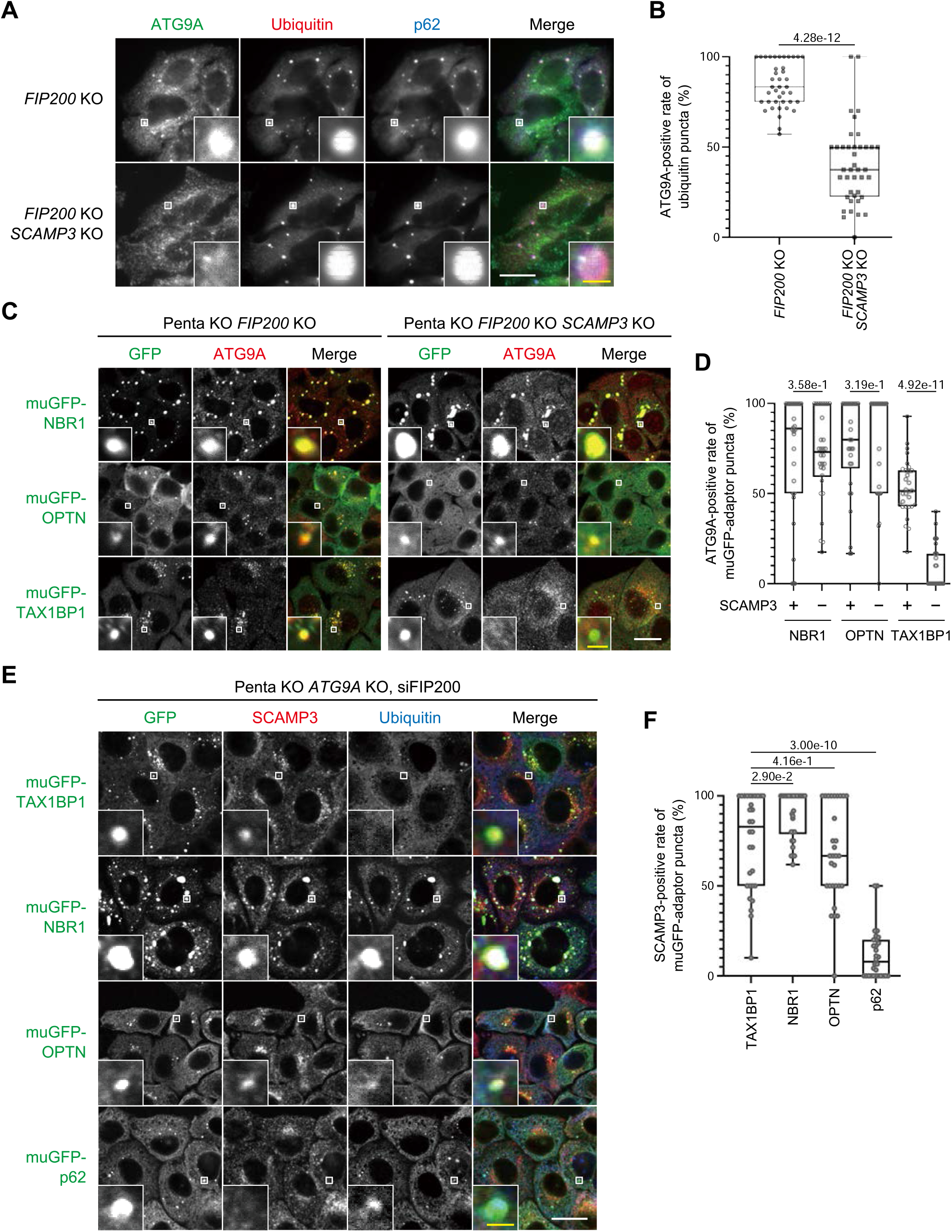
SCAMP3 is responsible for the TAX1BP1-dependent recruitment of ATG9 vesicles. **(A)** Recruitment of ATG9 vesicles to the p62 condensates is reduced in the absence of SCAMP3. *FIP200* KO and *FIP200* KO *SCAMP3* KO HeLa cells were immunostained with anti-ATG9A, anti-ubiquitin, and anti-p62 antibodies. **(B)** The ratios of ATG9-positive ubiquitin puncta to total ubiquitin puncta per cell in (A) were calculated (40 cells each) as in Figure 1D. **(C)** *SCAMP3* KO reduces recruitment of ATG9A to TAX1BP1, but not to NBR1 and OPTN. Penta KO *FIP200* KO and Penta KO *FIP200* KO *SCAMP3* KO HeLa cells stably expressing muGFP-tagged NBR1, OPTN, and TAX1BP1 were immunostained with anti-GFP and anti-ATG9A antibodies. **(D)** The ratios of ATG9-positive GFP puncta to total GFP puncta per cell in (C) were calculated (30 cells each) as in Figure 1D. **(E)** SCAMP3 localizes to the TAX1BP1 puncta even in the absence of ATG9A. Penta KO *FIP200* KO HeLa cells stably expressing muGFP-tagged TAX1BP1, NBR1, OPTN, and p62 were treated with siRNA against *FIP200*. Five days after the transfection, the cells were immunostained with anti-GFP and anti-ATG9A antibodies. **(F)** The ratios of ATG9-positive GFP puncta to total GFP puncta per cell in (E) were calculated (30 cells each) as in Figure 1D. White scale bars represent 20 μm and yellow scale bars represent 2 μm.

Additionally, we found that even in the absence of ATG9A, SCAMP3 localized to the muGFP-TAX1BP1 condensates in the Penta KO *FIP200* knockdown background (Fig. 6E, 6F), suggesting that ATG9A is not the determinant for the TAX1BP1- and SCAMP3-dependent recruitment of ATG9 vesicles. Furthermore, endogenous SCAMP3 is also localized to muGFP-NBR1 and muGFP-OPTN condensates in the absence of ATG9A (Fig. 6E, 6F). Taken together, we conclude that the TAX1BP1–SCAMP3 axis is responsible for the recruitment of ATG9 vesicles to the ubiquitin–p62 condensates independently of the ULK complex and other adaptors (Fig. 7).

**Figure 7.**
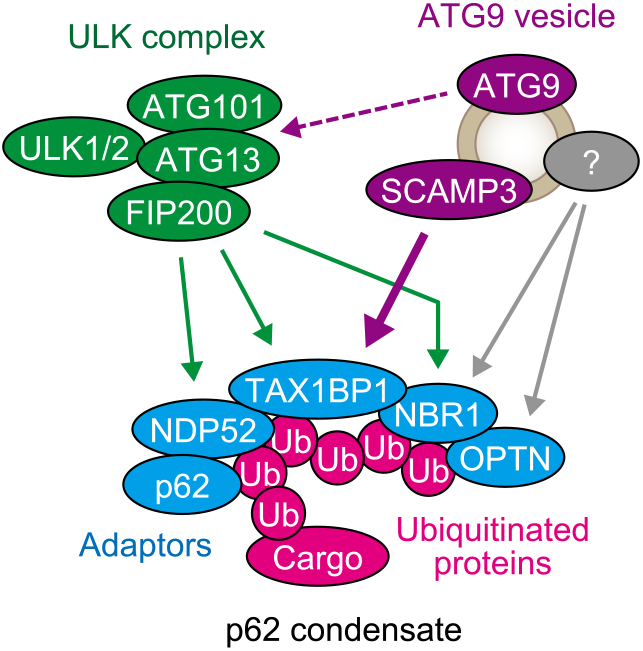
Cargo-dependent assembly of ATG9 vesicles dependent on the SCAMP3–TAX1BP1 interaction. ATG9 vesicles localize to ubiquitin-positive p62 condensates independently of the ULK complex, mediated by SCAMP3 present in ATG9 vesicles. The cargo-dependent assembly of ATG9 vesicles and the ULK complex are indicated by solid arrows, and the ULK complex-dependent recruitment of ATG9 vesicles is indicated by a dashed arrow.

## Discussion

Using the Penta KO cell line as a tool to study cargo-dependent recruitment of early ATG proteins, we found that ATG9 vesicles can be recruited by TAX1BP1, NBR1, or OPTN (Fig. 7). Of these, TAX1BP1, but not the other proteins, binds to SCAMP3 to recruit ATG9 vesicles (Fig. 5, 6). Furthermore, even in the absence of ATG9A, SCAMP3-containing structures were recruited to the TAX1BP1 condensates (Fig. 6). These observations indicate that SCAMP3 is the determinant for the TAX1BP1-dependent recruitment of ATG9 vesicles. Cargo-driven recruitment of early ATG proteins, such as ATG9 vesicles and the ULK complex, facilitates selective autophagy; the recruitment of FIP200 by the cargo adaptors p62, TAX1BP1, and NDP52 facilitates selective degradation of ubiquitin– p62 condensates and ubiquitinated mitochondria (Ravenhill et al. 2019; Turco et al. 2019; Vargas et al. 2019; Ohnstad et al. 2020; Turco et al. 2021), and the recruitment of ATG9 vesicles by OPTN leads to efficient mitophagy (Yamano et al. 2020). In this study, we sought to determine which process of selective autophagy is affected by deficiency of the TAX1BP1–SCAMP3–ATG9 vesicle axis. Whereas TAX1BP1 is known to be involved in the removal of ubiquitin–p62 condensates, ferritin–NCOA4 condensates, and intracellular pathogens (Tumbarello et al. 2015; Goodwin et al. 2017; Sarraf et al. 2020; Ohshima et al. 2022), we did not observe any obvious effects on the degradation of the ubiquitin–p62 condensates and ferritin–NCOA4 condensates in cultured cells. A previous study reported that TAX1BP1 is important for the removal of aggregates in mouse neurons (Sarraf et al. 2020). Thus, the recruitment of ATG9 vesicles by TAX1BP1 may contribute to the long-term effect of selective cargo degradation under physiological conditions. Another function of TAX1BP1 may also involve the TAX1BP1–SCAMP3–ATG9 vesicle axis, such as the selective degradation of the cytotoxic complex IIa formed upon tumor necrosis factor sensing (Huyghe et al. 2022). Future studies exploring the TAX1BP1–SCAMP3 axis in mice will provide further understanding of the physiological significance of TAX1BP1-dependent recruitment.

While NBR1 and OPTN can drive ATG9 vesicle translocation to ubiquitin– p62 condensates and ubiquitinated mitochondria, respectively (Sánchez-Martín et al. 2020; Yamano et al. 2020), the molecular details behind these cargo-dependent recruitments remain unclear. We showed that ATG9 vesicles were recruited to the NBR1 and OPTN condensates without SCAMP3 (Fig. 6). This finding implies that there are yet unknown molecules that mediate ATG9 vesicle recruitment by NBR1 and OPTN (Fig. 7). It is unclear why there are two or more distinct recruitment mechanisms for ATG9 – one by TAX1BP1 and the others by NBR1 and OPTN – but it is possible that these mechanisms are used differently for different cargos. Further analysis is needed to clarify this issue.

Previous studies have reported that ATG9A interacts with the ATG13– ATG101 subcomplex, which is involved in the recruitment of ATG9 vesicles to ubiquitin–p62 condensates and ubiquitinated mitochondria during selective autophagy (Kannangara et al. 2021; Ren et al. 2023). In this study, we showed that ATG9 vesicles are also recruited to the starvation-induced assembly of the ULK complex in a cargo/adaptor-independent manner (Fig. 2A, 2B). This finding indicates that the ULK complex contributes to the recruitment of ATG9 vesicles not only in selective autophagy but also in starvation-induced autophagy. This ATG9 vesicle recruitment is quite similar to that in yeast; upon starvation, yeast Atg9 vesicles are recruited to the preautophagosomal structure formed by the Atg1 complex through Atg9 binding to Atg13^HORMA^ (Suzuki et al. 2015). In addition, yeast Atg9 vesicles are also recruited to selective cargos via the adaptor Atg11, which may be analogous to the cargo/adaptor-dependent recruitment of mammalian ATG9 vesicles. These findings suggest that two types of ATG9 vesicle recruitment mechanisms, namely the cargo/adaptor-dependent assembly and ULK/Atg1-dependent assembly, have been conserved during evolution.

## Acknowledgments

We would like to thank Michael Lazarou for providing the Penta KO HeLa cells; Shoji Yamaoka for the pMRXIP plasmid; and Teruhito Yasui for the pCG-gag-pol and pCG-VSV-G plasmids. This work was supported by a Grant-in-Aid for Transformative Research Areas (A) (21H05256 to H.Y.) and for Specially Promoted Research (22H04919 to N.M.) from the Japan Society for the Promotion of Science (JSPS) and the Exploratory Research for Advanced Technology (ERATO) research funding program of the Japan Science and Technology Agency (JST) (JPMJER1702 to N.M.).

## Author Contributions

Y.H., N.M., and H.Y. designed the project. Y.H. and T.M. carried out the cell biology experiments. Y.K. performed the mass spectrometry. Y.H., N.M., and H.Y. wrote the manuscript. All authors have commented on the manuscript.

## Materials and Methods

### Cell culture

HeLa and HEK293T cells (authenticated by RIKEN) were cultured in Dulbecco’s Modified Eagle medium (DMEM) (043-30085; Fujifilm Wako Pure Chemical Corporation) supplemented with 10% fetal bovine serum (FBS) (172012; Sigma-Aldrich) in a 5% CO_2_ incubator at 37°C. For starvation, cells were washed twice with phosphate-buffered saline (PBS) and incubated in amino acid-free DMEM (048-33575; Fujifilm Wako Pure Chemical Corporation) without FBS.

### Plasmids

Plasmids for deletion of the *SCAMP3*, *FIP200*, *ATG9A*, and *ATG13* genes were generated as follows: sgRNA targeting *SCAMP3* (5′-GTGGGGCTGAGCTTTCTCGA-3′), *FIP200* (5′-TATGTATTTCTGGTTAACAC-3′), *ATG9A* (5′-CCTGTTGGTGCACGTCGCCG-3′), and *ATG13* (5′-CCGCGAGTTTGATGCCTTTG-3′) were inserted into the BpiI site of PX458 [pSpCas9(BB)-2A-GFP; Addgene #48138].

Plasmids for stable expression in HeLa cells were generated as follows: DNA fragments encoding human SCAMP3 (NM_005698.4), ATG13 (Hosokawa et al. 2009), ATG101 (Hosokawa et al. 2009), ATG9A (Nishimura et al. 2017), p62, NBR1 (Itakura and Mizushima 2011), OPTN (NM_001008211.1), NDP52 (NM_001261390), and TAX1BP1 (isoform 2, NM_001079864) were inserted into the retroviral plasmid pMRX-IP (puromycin-resistant marker) (Kitamura et al. 2003; Saitoh et al. 2003), pMRX-IB (blasticidin-resistant marker) (Morita et al. 2018), or pMRX-IH (hygromycin-resistant marker) together with muGFP, mRFP, or 3×FLAG. pMRX-IH was constructed by replacing the puromycin-resistant gene of pMRX-IP with the hygromycin-resistant gene.

### Generation of knockout cell lines

*FIP200* KO, *ATG9A* KO (Tsuboyama et al. 2016), and Penta KO (Lazarou et al. 2015) HeLa cell lines have been described previously. For further deletion of the *SCAMP3*, *ATG13*, *FIP200*, or *ATG9A* gene, HeLa cells were transfected with the PX458-based plasmid expressing Cas9-T2A-GFP with sgRNA for the *SCAMP3* gene, the *ATG13* gene, the *FIP200* gene, or the *ATG9A* gene using ViaFect (E4981; Promega). Two days after transfection, GFP-positive cells were isolated using a cell sorter (MoFlo Astrios EQ; Beckman Coulter) and single clones were obtained.

### Stable expression in HeLa cells by retroviral infection

HEK293T cells were transfected with the pMRX-IP-, pMRX-IB-, or pMRX-IH-based retroviral plasmid (Kitamura et al. 2003; Saitoh et al. 2003) together with pCG-gag-pol and pCG-VSV-G (a gift from Dr. T. Yasui, National Institutes of Biomedical Innovation, Health and Nutrition) using FuGENE HD (E2311; Promega) for 4 h in Opti-MEM (31985-070; Gibco). After cultivating the cells for 2–3 days in DMEM, the retrovirus-containing medium was harvested, filtered through a 0.45-µm filter unit (Ultrafree-MC; Millipore), and added to HeLa cells with 8 μg/ml polybrene (H9268; Sigma-Aldrich). After cultivating the cells for 24 h, stable transformants were selected by treatment with 2 μg/ml puromycin (P8833; Sigma-Aldrich), 3 μg/ml blasticidin (022-18713; Fujifilm Wako Pure Chemical Corporation), or 50 μg/ml hygromycin (10687010; Thermo Fisher Scientific).

### RNA interference

Stealth RNAi siRNAs (Thermo Fisher Scientific) were used for RNA interference. Penta KO *ATG9A* KO HeLa cells were transfected with siRNA against *FIP200* (GGAAAUGUAUGAAGUUGCCAAGAAA) using Lipofectamine RNAiMAX (13778150; Thermo Fisher Scientific) for 4 h in Opti-MEM (31985-070; Gibco). The cells were cultured in DMEM for 56 h.

### Preparation of whole cell lysates and immunoblotting

HeLa cells were lysed with 0.2% n-octyl-β-D-dodecyl maltoside (DDM) in lysis buffer [50 mM Tris-HCl (pH 7.5), 150 mM NaCl, 1 mM EDTA, cOmplete EDTA-free protease inhibitor cocktail (05056489001; Roche)] for 15 min on ice. After centrifugation at 17,700 × *g* for 15 min at 4°C, the supernatants were collected and mixed with 20% volumes of 6×SDS-PAGE sample buffer. The samples were subjected to SDS-PAGE and transferred to Immobilon-P PVDF membranes (IPVH00010; EMD Millipore). Immunoblotting was performed using rabbit polyclonal antibodies against SCAMP3 (26888-1-AP; ProteinTech), ATG9A (PD042; MBL), FIP200 (17250-1-AP; ProteinTech), RAB1A (sc-311; Santa Cruz), and RCAS1 (12290T; CST) and mouse monoclonal antibody against HSP90 (610419; BD Biosciences) as primary antibodies, and horseradish peroxidase (HRP)-conjugated anti-rabbit IgG (111-035-144; Jackson ImmunoResearch) as a secondary antibody. Super-Signal West Pico Chemiluminescent Substrate (1856135; Thermo Fisher Scientific) and Immobilon Western Chemiluminescent HRP Substrate (P90715; EMD Millipore) were used to visualize the signals, which were detected by an image analyzer (FUSION SOLO.7S.EDGE; Vilber-Lourmat). Contrast and brightness adjustment and quantification were performed using Fiji software (ImageJ; National Institutes of Health) (Schindelin et al. 2012).

### Immunostaining and fluorescence microscopy

HeLa cells were grown on coverslips, washed with PBS, and fixed with 4% paraformaldehyde (09154-85; Nacalai Tesque) in PBS for 10 min on ice. After permeabilization with 50 µg/ml digitonin in PBS for 5 min and blocking with 3% BSA in PBS for 30 min at room temperature, the cells were incubated with primary antibodies in PBS containing 3% BSA for 1 h at room temperature, washed five times with PBS, incubated with secondary antibodies in PBS containing 3% BSA for 1 h at room temperature, and washed five times with PBS. The cells were mounted with SlowFade Gold Antifade Mountant with DAPI (S36939; Thermo Fisher Scientific) and observed with a confocal microscope (FV1000; Olympus).

Immunostaining was performed using rabbit polyclonal antibodies against ATG9A (ab108338; abcam), FIP200 (17250-1-AP; ProteinTech), and SCAMP3 (SAB2701979; Sigma-Aldrich), mouse monoclonal antibodies against FLAG (F1804; Sigma-Aldrich) and ubiquitin (D058-3; MBL), rat monoclonal antibody against GFP (D153-3; MBL), and guinea pig polyclonal antibodies against p62 (GP62-C; PROGEN) as primary antibodies. Alexa Fluor 488-conjugated anti-mouse IgG (also cross-adsorbed to rabbit IgG and rat IgG) (A-11029; Thermo Fisher Scientific) and Alexa Fluor 568-conjugated anti-rabbit IgG (also cross-adsorbed to mouse IgG) (A-11036; Thermo Fisher Scientific) were used as secondary antibodies.

### Sample preparation for LC–MS/MS analysis

Penta KO *FIP200* KO HeLa cells stably expressing FLAG-TAX1BP1^CC1^ (WT and K248E) and Penta KO *FIP200* KO HeLa cells (as a non-expressing control) were solubilized with 0.2% DDM in lysis buffer for 15 min on ice. After centrifugation at 17,700 × *g* for 15 min at 4°C, the supernatants were collected and incubated with anti-FLAG M2 affinity gel (A2220; Sigma-Aldrich) for 3 h at 4°C with gentle rotation. The beads were precipitated at 700 × *g* for 10 sec at 4°C, washed three times with 0.05% DDM in wash buffer [20 mM Tris-HCl (pH 8.0), 150 mM NaCl], and immunoprecipitates were eluted with 100 µg/ml 3×FLAG peptide (4046200; Bachem AG) in wash buffer. Subsequent sample preparation for LC–MS/MS experiments was carried out according to the phase-transfer surfactant protocol (Masuda et al. 2008). Samples were reduced with 10 mM dithiothreitol and alkylated with 50 mM 2-iodoacetamide (804744; Sigma-Aldrich), followed by digestion with Lysyl Endopeptidase (121-05063; Fujifilm Wako Pure Chemical Corporation) and trypsin at 37°C overnight. The digest was acidified with 0.5% trifluoroacetic acid (TFA), and sodium deoxycholate was extracted with an equal volume of ethyl acetate and vigorous shaking. The organic phase was removed after centrifugation at 10,000 × *g* for 10 min at room temperature. The aqueous phase containing the peptides was collected and dried using a centrifugal evaporator. The dried peptides were solubilized in 100 µl of 2% acetonitrile and 0.1% TFA, and the peptide mixture was captured on a handmade C18 STAGE tip prepared as previously reported (Rappsilber et al. 2007). The trapped peptides were subjected to dimethyl labeling with formaldehyde as previously described (Boersema et al., 2009). Briefly, the peptides prepared from FLAG-TAX1BP1^CC1^ (WT) cells, FLAG-TAX1BP1^CC1^ (K248E) cells, and non-expressing cells were heavy, medium, and light dimethyl-labeled with isotope-labeled formaldehyde (^13^CD_2_O; ISOTEC), isotope-labeled formaldehyde (CD_2_O; CIL), and formaldehyde (CH_2_O; SIGMA), respectively. The dimethyl-labeled peptides remaining on the tip were eluted with 100 μl of 80% acetonitrile and 0.1% TFA. All the eluates from the same species were mixed and dried using a centrifugal evaporator.

### Liquid chromatography–tandem mass spectrometry

The LC–MS/MS experiments were performed using a nanoscale liquid chromatography system (nanoLC-Ultra; Ekisgent) equipped with an Ekspert cHiPLC system (Eksigent) connected to a TripleTOF 5600+ mass spectrometer (AB Sciex). The sample acquisitions were performed using a ‘trap and elute’ configuration on the nanoFlex system. The trap column (200 µm × 0.5 mm) and an analytical column (75 µm × 15 cm) were packed with 3 µm ChromXP C18 medium. The dried peptide samples were dissolved in 15 µl of 2% acetonitrile with 0.1% TFA and subsequently loaded to the LC–MS system to be separated by a 2– 40% linear gradient of acetonitrile containing 0.1% formic acid for 200 min at a flow rate of 300 nl/min. The eluted peptides were electrosprayed (2.3 kV) and injected into the mass spectrometer in the data-dependent acquisition mode. Precursor ion spectra were acquired over the 400–1,250 *m/z* mass range, and the top 20 most intense ions were selected for MS/MS analyses.

### Peptide identification and quantification

All MS/MS spectra were searched against protein sequences in the NCBI protein dataset (NCBInr RefSeq Release 90) using the Protein Pilot software package (AB Sciex).

### Isolation of ATG9 vesicles

Cell homogenates were prepared by 10 passages through a 27-gauge syringe at 4°C in 25 mM HEPES, 100 mM sorbitol, 5 mM EDTA, 100 mM urea, 100 mM arginine, 50 mM NaCl, and 0.5 mg/ml BSA. The homogenates were centrifuged at 1,000 × *g* for 10 min at 4°C and the supernatants were ultracentrifuged at 50,000 × *g* for 30 min at 4°C. The supernatants (input) were incubated with anti-FLAG M2 affinity gel (A2220; Sigma-Aldrich) for 3 h at 4°C with gentle rotation. The affinity gels were precipitated at 700 × *g* for 10 sec at 4°C and washed three times, and immunoprecipitates were eluted with 100 µg/ml 3×FLAG peptide (4046200; BACHEM).

### Statistical analysis

Statistical analyses were performed using GraphPad Prism 9 software (GraphPad Software). The statistical methods are described in each figure legend.

---

**Supplementary Figure S1.**
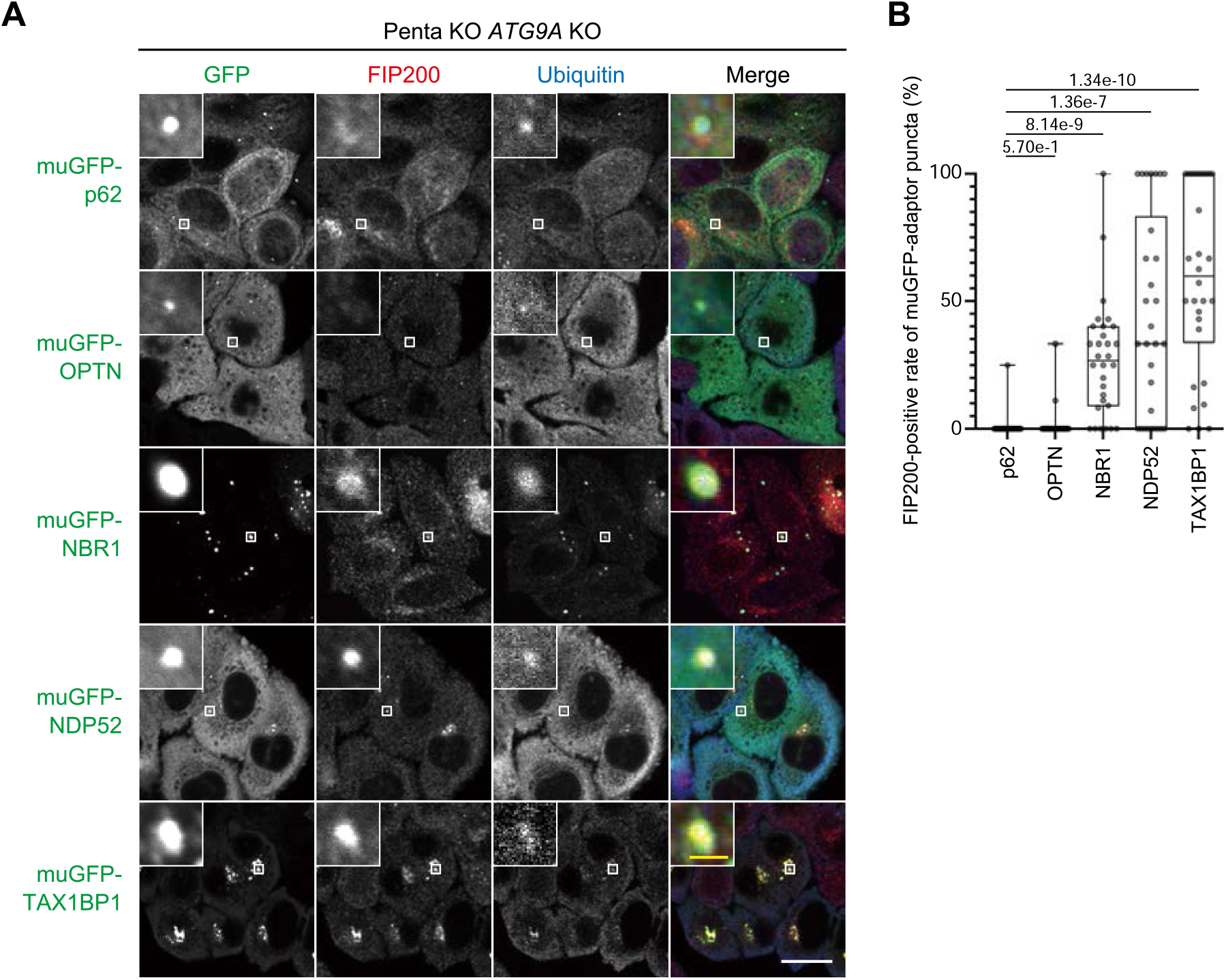
NBR1, NDP52, and TAX1BP1 independently recruit the ULK complex. **(A)** The ULK complex is recruited by NBR1, NDP52, and TAX1BP1. Penta KO *ATG9A* KO HeLa cells stably expressing muGFP-tagged p62, OPTN, NBR1, NDP52, or TAX1BP1 were immunostained with anti-GFP, anti-FIP200, and anti-ubiquitin antibodies. **(B)** The ratios of FIP200-positive GFP puncta to total GFP puncta per cell in (A) were calculated (30 cells each) as in Figure 1D. The white scale bar represents 20 μm, and the yellow scale bar represents 2 μm.

**Supplementary Figure S2.**
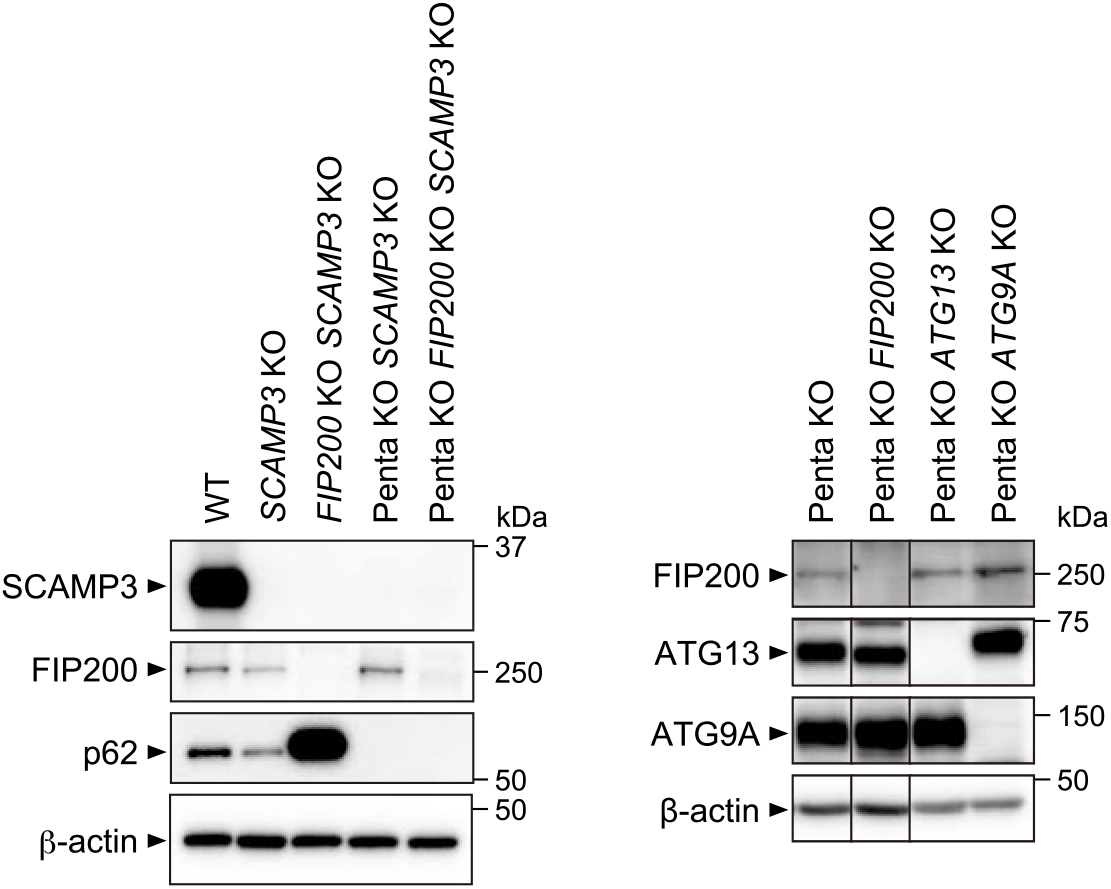
Immunoblotting of *SCAMP3* KO, *FIP200* KO, *ATG13* KO, and *ATG9A* KO cells. WT, *SCAMP3* KO, *FIP200* KO *SCAMP3* KO, Penta KO *SCAMP3* KO, Penta KO *FIP200* KO *SCAMP3* KO, Penta KO, Penta KO *FIP200* KO, Penta KO *ATG13* KO, and Penta KO *ATG9A* KO HeLa cells were grown in DMEM. Whole-cell lysates were analyzed by immunoblotting with antibodies against SCAMP3, FIP200, ATG13, ATG9A, p62, and β-actin.

**Supplementary Table S1.** LC–MS/MS results of the TAX1BP1^CC1^ interactome. Penta KO *FIP200* KO HeLa cells stably expressing FLAG-TAX1BP1^CC1^ (WT or K248E) and Penta KO *FIP200* KO HeLa cells as non-expressing cells were solubilized with 0.2% DDM and subjected to immunoprecipitation with anti-FLAG beads. The immunoprecipitates were labeled by dimethyl labeling to FLAG-TAX1BP1^CC1^ (WT) as heavy (H), FLAG-TAX1BP1 (K248E) as medium (M), and non-expression of TAX1BP1 as light (L) and analyzed by LC–MS/MS. The ratios of FLAG-TAX1BP1^CC1^ WT to non-expression (H/L) and FLAG-TAX1BP1^CC1^ K248E to non-expression (M/L) are shown with the order of the H/L ratio.

